# Minimal Tool Set for a Prokaryotic Circadian Clock

**DOI:** 10.1101/075291

**Authors:** Nicolas M Schmelling, Robert Lehmann, Paushali Chaudhury, Christian Beck, Sonja V Albers, Ilka M Axmann, Anika Wiegard

## Abstract

**Background:** Circadian clocks are found in organisms of almost all domains including photosynthetic Cyanobacteria, whereby large diversity exists within the protein components involved. In the model cyanobacterium *Synechococcus elongatus* PCC 7942 circadian rhythms are driven by a unique KaiABC protein clock, which is embedded in a network of input and output factors. Homologous proteins to the KaiABC clock have been observed in Bacteria and Archaea, where evidence for circadian behavior in these domains is accumulating. However, interaction and function of non-cyanobacterial Kai-proteins as well as homologous input and output components remain mainly unclear.

**Results:** Using a universal BLAST analyses, we identified putative KaiC-based timing systems in organisms outside as well as variations within Cyanobacteria. A systematic analyses of publicly available microarray data elucidated interesting variations in circadian gene expression between different cyanobacterial strains, which might be correlated to the diversity of genome encoded clock components. Based on statistical analyses of co-occurrences of the clock components homologous to *Synechococcus elongatus* PCC 7942, we propose putative networks of reduced and fully functional clock systems. Further, we studied KaiC sequence conservation to determine functionally important regions of diverged KaiC homologs. Biochemical characterization of exemplary cyanobacterial KaiC proteins as well as homologs from two thermophilic Archaea demonstrated that kinase activity is always present. However, a KaiA-mediated phosphorylation is only detectable in KaiC1 orthologs.

**Conclusion:** Our analysis of 11,264 genomes clearly demonstrates that components of the *Synechococcus elongatus* PCC 7942 circadian clock are present in Bacteria and Archaea. However, all components are less abundant in other organisms than Cyanobacteria and KaiA, Pex, LdpA, and CdpA are only present in the latter. Thus, only reduced KaiBC-based or even simpler, solely KaiC-based timing systems might exist outside of the cyanobacterial phylum, which might be capable of driving diurnal oscillations.

## Introduction

Life on Earth is under the influence of changing environmental conditions, which not only pose a challenge to organisms, but also present a chance of adaptation and therefore a possible fitness advantage over competitors [1, 2]. Using inner timing systems organisms can coordinate their physiology and behavior according to the daily recurring changes. Simple timing systems work in an hour glass like fashion and need to be reset every day by environmental stimuli, whereas true circadian clocks generate self-sustained and temperature-compensated 24-hour rhythms of biological activities [3, 4].

Circadian clocks are found in many eukaryotes such as algae, plants and mammals [5]. Even though circadian clocks seem like a conserved trait in evolution, differences in the protein components, involved in circadian timing, suggest a convergent evolution of timing mechanisms [5]. For many years it has been believed that something as complex as a circadian clock could not have been evolved in unicellular organisms like prokaryotes [5, 6, 7]. However, the existence of temperature compensated 24-hour rhythms of cell division in *Synechococcus* sp. WH 7803 and circadian nitrogen fixation in *Cyanothece* sp. PCC 8801 proved otherwise [8, 9, 10, 11, 12]. The molecular basis of the cyanobacterial circadian clock was intensively investigated in *Synechococcus elongatus* PCC 7942 (hereafter *Synechococcus* 7942), where the core clockwork resembles a posttranslational oscillator [13, 14, 15]. In contrast, eukaryotic circadian rhythms are believed to be mainly based on transcriptional-translational feedback loops. However, findings on post-translational systems are accumulating [16, 17, 18] and might exist also in Archaea [16].

Light is assumed to be the driving stimulus in circadian clock entrainment [19]. In *Synechococcus* 7942, contrary to eukaryotic circadian clock systems, a photoreceptor in the input pathway of the clock could not be detected thus far. Instead, *Synechococcus* 7942 cells sense light indirectly through the redox and energy state of the cell [20]. Here, two metabolic components are considered to play a major role [21]: The ATP to ADP ratio and the redox state of the plastoquinone (PQ) pool [22, 23]. The core of the circadian clock in *Synechococcus* 7942 consists of three proteins KaiA, KaiB and KaiC. KaiC monomers are composed of two domains, which assemble into two hexameric rings [24, 25]. The C-terminal ring is capable of autophosphorylation and –dephosphorylation [26, 27]. KaiC phosphorylation is stimulated by the interaction with KaiA [28, 29], and additionally, affected by the ATP/ADP ratio of the cell [30]. KaiB inhibits the activating effect of KaiA and initializes dephosphorylation [31]. Altogether, KaiC hexamers phosphorylate and dephosphorylate rhythmically during the course of a day. The binding of oxidized quinones to KaiA has been suggested to stop the clock directly by causing KaiA aggregation [32, 33]. The KaiABC core clock is embedded into a network of input and output pathways. The input factors that interact with the core clock are Pex, LdpA, PrkE, NhtA, IrcA, CdpA [20, 34, 35, 36, 37, 38]. Output factors are SasA, LabA, LalA, CpmA, Crm, RpaA, and RpaB, as well as CikA, which is functioning both in input and output pathway of the circadian clock [39, 40, 41, 42, 43, 44, 45, 46, 47, 48, 49, 50, 51].

Sequence analysis indicated that at least three different types of timing systems are present in Cyanobacteria, (i) a KaiABC-based system as in *Synechococcus* 7942, (ii) a reduced system with a KaiBC core and a reduced set of input/output factors as in *Prochlorococcus* and (iii) a reduced KaiABC system as in *Synechococcus* sp. WH 8102, which despite including all three *kai* genes, has the same input/output factors as the reduced KaiBC system [52]. Furthermore, multiple *kai* genes can exist in an organism [53, 54]. In *Synechocystis* sp. PCC 6803 (hereafter *Synechocystis* 6803) for example, three copies of both *kaiB* and *kaiC* are found. KaiA, KaiB1 and KaiC1, most similar to the *kai* genes of *Synechococcus* 7942 [55, 56], are thought to resemble the core clock. KaiB3 and KaiC3 are thought to function as fine tuning factors for the circadian clock in *Synechocystis* 6803, whereas no circadian function has been found for KaiB2 and KaiC2 [56]. However, recently *kaiC2*-dependent adaptive growth and diurnal rhythms of nitrogen fixation were observed in *Rhodopseudomonas palustris* [57]. Homologs of *kaiB* and *kaiC* genes also exist in other Bacteria and even Archaea, where a shortend KaiC most often resembles only one domain [53]. An archaeal one-domain KaiC homolog was shown to form a hexameric ring, similar to the two duplicated domains of *Synechococcus* 7942 KaiC [58]. In *Haloferax volcanii* the transcripts of four *kaiC* homologs display diurnal accumulation profiles [59]. However, the function of non-cyanobacterial Kai-proteins is mainly unclear, so far. Although some of the input- and output factors were also found in prokaryotes other than cyanobacteria [49, 52, 60, 61], it is unknown whether the Kai homologs outside the cyanobacterial phylum or the additional cyanobacterial Kai homologs interact with (other) in- and output factors.

In this study, we performed BLAST analyses to first identify possible KaiC-based timing systems in organisms outside of Cyanobacteria and second to explore variations in circadian clocks within Cyanobacteria. Further, we examined variations in circadian gene expression between different cyanobacterial strains using microarray data. Together, this aims at decoding the correlation between Kai proteins and additional clock components. Based on the co-occurrence of clock components known from *Synechococcus* 7942 we propose putative networks of reduced and fully functional clock systems. Further, we used the sequence information of KaiC and its homologs in Cyanobacteria to determine the similarities at important sites of the protein. We chose cyanobacterial KaiC proteins as well as homologs from two thermophilic Archaea and demonstrated that kinase activity is always present. However, a KaiA-mediated phosphorylation is only detectable in cyanobacterial KaiC1 homologs.

## Materials and Methods

### Programming languages

The programming languages Python (version 3.5.1) and R (version 3.2.3) were used in this work. The processing and analysis of the microarray time series datasets was performed using R. Regarding the distribution analysis, the Biopython project ([62]; version 1.66) was used to download from GenBank as well as to work with FASTA files. Besides Biopython, the Python packages: IPython ([63] version 4.1.1) as an interactive Python environment with the IPython notebook; numpy and scipy ([64]; version 1.10.4, version 0.17.0) for numerical operations; matplotlib ([65]; version 1.5.1) for data visualization; and pandas ([66]; version 0.17.1) for data analyses were used. The Python code necessary to reproduce the BLAST analyses is available on GitHub (http://doi.org/10.5281/zenodo.229910).

### Reciprocal BLAST and NCBI

The coding sequences of all entries in the genbank protein database [67], which were labeled as “Complete Genome” or “Chromosome”, were downloaded from the NCBI FTP server (version May 2016). These sequences, including the coding sequences of *Synechococcus elongatus* PCC 7942 and *Synechocystis* sp. PCC 6803, were used to construct a custom protein database for the homology search. Further, protein sequences of the 23 clock related proteins (Table S1), from *Synechococcus elongatus* PCC 7942 and *Synechocystis* sp. PCC 6803, respectively, were checked against the entries in the Cyanobase Database to ensure correctness [68] (version May 2016). These 23 protein sequences were used as queries for a search of homologs within the custom protein database, applying the standalone version of BLASTP 2.2.30+ [69] (May 2016, dx.doi.org/10.17504/protocols.io.grnbv5e) with standard parameter (wordsize: 3, substitution matrix: BLOSUM62). The 10,000 best hits with an e-value of 10*^−^*^5^ or lower were filtered for further analyses. The first BLAST run returned circa 65,000 hits for all 23 cyanobacterial proteins combined.

These hits were used as queries for a second reverse BLASTP run, searching for homologs in *Synechococcus* 7942 or *Synechocystis* 6803 genomes using the same parameters as above with an altered e-value of 10. Only hits with the original query protein as best reversal hit were accepted for further analyses, thus minimizing false positive results.

Raw and processed data is available on figshare (https://dx.doi.org/10.6084/m9.figshare.3823902.v3, https://dx.doi.org/10.6084/m9.figshare.3823899.v3).

### Testing of co-occurence

Co-occurrence of circadian clock proteins was examined by using the right-sided Fisher’s exact test [70]. For each of the 94 cyanobacterial strains, all identified homologous clock genes were gathered into one set. The phylogenetic distribution of cyanobacteria in the NCBI genbank database is very imbalanced. Some genera (e.g. *Prochlorococcus* and *Synechococcus*) are covered better than others. To avoid selection bias, we removed sets with identical combinations of genes, resulting in 69 unique clock systems. Null hypothesis of Fisher’s exact test is a pairwise independent distribution of the proteins across all clock systems. P-values were corrected for multiple testing after Benjamini-Hochberg [71] with an excepted false discovery rate of 10*^−^*^2^. We denote that due to the nature of statistical testing, proteins appearing in almost all clock systems are always virtually independent to others. All proteins were clustered according to their corrected p-values.

### Microarray analysis

The diurnal expression program of six cyanobacterial strains was probed using microarray time series datasets (Table S3). Unfortunately, the data for the two reported *Synechocystis* 6803 experiments by Labiosa and colleagues [72] and Kucho and coworkers [73] could not be obtained. The study by Toepel and coworkers [74] had to be discarded due to the employed ultradian light cycles (6:6 LD cycles). The *Synechocystis* 6803 datasets, the *Synechococcus* 7942 dataset of Ito and colleagues, and the *Anabaena* sp. PCC 7120 dataset were l2m transformed, while the *Cyanothece* ATCC 51142 datasets are only available after transformation. The two biological replicates of the *Anabaena* sp. PCC 7120 dataset were concatenated for the following analyses [75], similar to the *Synechocystis* 6803 dataset [76]. Expression profiles were smoothed using a Savitzky-Golay lowpass filter, as proposed by Yang and colleagues [77], in order to remove pseudo peaks prior to the detection of periodic genes. Diurnally oscillating expression profiles were detected using harmonic regression analysis [78]. The derived p-values for each gene is based on the assumption of a linear background profile as compared to the sinoidal foreground model. After multiple hypothesis testing correction according to Benjamini-Hochberg [71], all datasets yielded significantly oscillating genes (q ≤ 0.05) except for *Microcystis aeruginosa* PCC 7806, with 7 samples the shortest dataset.

### Multiple sequence alignment

Multiple sequence alignments were constructed by the standalone version of CLUSTAL Omega 1.2.1-1 [79] using 20 iterations, while only one iteration was used to construct the guide tree (May 2016, dx.doi.org/10.17504/protocols.io.gscbwaw). The sequences for the alignments were obtained from the processed data generated, as described in the method section “Reciprocal BLAST and NCBI”. Afterwards the alignments were adjusted to *Synechococcus* 7942 sequence with Jalview [80] and edited multialignments were used to create WebLogos [81].

### Cloning, heterologous expression and purification of Kai proteins

To express KaiC1 proteins from *Synechocystis* sp. PCC 6714, *Nostoc punctiforme* ATCC 29133, *Cyanothece* sp. PCC 7424 as well as KaiC3 from *Cyanothece* sp. PCC 7424, *Microcystis aeruginosa* PCC 7806, *Pycrococcus horikoshii* OT3 PH0833, and *Thermococcus litoralis* DSM5473, the respective *kaiC* genes were amplified by PCR from genomic wildtype DNA. The ORFs and primers are listed in Table S4. Amplified sequences were ligated into BamHI and NotI restriction sites of the plasmid pGEX-6P-1 (GE Healthcare). For *P.horikoshii* and *T.litoralis* KaiC3 the amplified PCR products were ligated into pETDuet1 Vector using BamHI/HindIII and PstI/HindIII restriction enzymes in MCS1 respectively. *Escherichia coli* DH5*α* or BL21 (DE3) cells were transformed with the resulting plasmids (pGEX-kaiC1-*Sy* 6714, pGEX-kaiC1-*Npun*29133, pGEX-kaiC1-*Cy*7424, pGEX-kaiC3-*Cy*7424, pGEX-kaiC3-*Mic*7806, pSVA3151, pSVA3152). For expression of GST-fused KaiA-7942 and KaiC-7942 pGEX derivatives, kindly provided by T. Kondo (Nagoya University, Japan), were used. Expression of GST-Kai proteins occurred at 37 *^◦^*C and 200 rpm in Terrific broth medium containing 100 *µ*g ampicillin ml*^−^*^1^. GST-KaiA-7942 expression was induced with 1 mM isopropyl *β*-D-thiogalactopyranoside (IPTG) and carried out overnight, whereas GST-KaiC homologs were expressed for 72 hours without induction. Cells were harvested and lysed in ice-cold extraction buffer (50 mM Tris/HCl (pH 8), 150 mM NaCl, 0.5 mM EDTA, 1 mM DTT, 5 mM MgCl_2_ and 1 mM ATP). Recombinant GST-Kai proteins were affinity purified using Protino Gluthatione Agarose 4B (Macherey–Nagel) as described in Wiegard and colleagues [55]. During the procedure the GST-tag was removed with PreScission protease in cleavage buffer (50 mM Tris/HCl (pH 8), 150 mM NaCl, 1 mM EDTA, 1 mM DTT, 5 mM MgCl_2_ and 0.5 mM ATP). Homogeneity of the recombinant proteins was controlled by separating them via SDS-PAGE. If it was not sufficient, proteins were further purified by anion-exchange chromatography using a MonoQ 5/50 GL or ResourceQ column (GE Healthcare). After dialysis in reaction buffer (20 mM Tris/HCl (pH 8), 150 mM NaCl, 0.5 mM EDTA, 5 mM MgCl_2_, and 1 mM ATP) protein concentration was determined using infrared spectroscopy (Direct detect, Merck Millipore). Proteins were stored at -20 *^◦^*C. For His-tagged KaiC3 homologs, *E. coli* BL21 (DE3) RIL cells were transformed with pSVA3151 (*P.horikoshii*) and pSVA3152 (*T.litoralis*), and grown as preculture overnight at 37 *^◦^*C in LB medium containing ampicillin (50 *µ*g ml*^−^*^1^) and chloramphenicol (34 *µ*g ml*^−^*^1^). Fresh medium containing antibiotic was inoculated with 0.1 % preculture and grown at 37 *^◦^*C to an OD_600_ of 0.7. After induction with 0.3 mM of IPTG, growth was continued for 16 hours at 16 *^◦^*C. Cells were collected by centrifugation, frozen in liquid nitrogen, and, after storage at -80 *^◦^*C, resuspended in 50 ml lysis buffer (50 mM Hepes-NaOH, pH 7.2, 150 mM NaCl) containing Complete EDTA-free protease inhibitor cocktail (Roche) together with DNase I and lysed by sonication. Cell debris were removed by centrifugation at 4 *^◦^*C for 30 min at 20,000 x g. Further Ni-NTA (Sigma Aldrich, Seelze, Germany) based purification was performed using columns equilibrated in purification buffer (50 mM Hepes-NaOH, pH 7.2, 150 mM NaCl). For the removal of unspecifically bound protein columns were washed with 15 column volumes of equilibration buffer including 10 mM imidazole. *T.litoralis* KaiC3 was eluted in the same buffer with 150 mM imidazole containing equilibration buffer, whereas for *P.horikoshii* KaiC3 the elution was carried out in 20 mM MES pH 6.2, 150 mM NaCl, 150 mM imidazole as this protein is stable in low pH buffer. Further purification of *T.litoralis* KaiC3 was achieved by size exclusion chromatography using Superdex 200 10/300 GL. *P.horikoshii* KaiC3 was incubated at 50 *^◦^*C for 20 min and centrifuged for 15 min at 10.000 x g. Subsequently, the supernatant was dialyzed overnight against 20 mM MES pH 6.2, 150 mM NaCl buffer. As a quality control, proteins were separated via SDS-PAGE and pure proteins were frozen in liquid nitrogen and kept at -80 *^◦^*C.

### *In vitro* phosphorylation assays

To investigate KaiA dependent phosphate uptake 12 *µ*g of KaiC-7942, KaiC1-*Sy*6714, KaiC1-*N*29133, KaiC1-*Cy*7424, KaiC3-*Cy*7424, KaiC3-*Mic*7806, KaiC3-*T.lit* or KaiC3-*P.hor* were mixed with 10 *µ*Ci *γ*-P^32^-ATP in 60 *µ*l Tris reaction buffer (20 mM Tris/HCl (pH 8), 150 mM NaCl, 0.5 mM EDTA, 5 mM MgCl_2_, 1 mM ATP) in the presence or absence of 6 *µ*g KaiA-7942. 10 *µ*l aliquots were taken after 0, 0.75, 1.5, 3 and 22 hours of incubation at 30 *^◦^*C and reaction was stopped by adding SDS-sample buffer. Proteins were separated in high-resolution polyacrylamide gels (10 % T, 0.67 % C) by SDS-PAGE (modified from [82]), stained with Coomassie brilliant blue and subjected to autoradiography. Signals were analyzed using a Fujifilm FLA-3000 (FUJIFILM). To analyze *in vitro* phosphorylation of KaiC3-*T.lit* and KaiC3-*P.hor* at higher temperatures, 10 *µ*g of the recombinant proteins were incubated with 10 *μ*Ci *γ*-P^32^-ATP in 50 *µ*l HEPES reaction buffer (50 mM HEPES (pH 7.2), 150 mM NaCl, 5 mM MgCl_2_, 1 mM ATP) or 50 *µ*l MES reaction buffer (50 mM MES (pH 6), 150 mM NaCl, 5 mM MgCl_2_, 1 mM ATP), respectively, at 75 *^◦^*C. After 0, 5, 10 and 15 minutes 10 *µ*l aliquots were taken and analyzed by SDS-PAGE and autoradiograpy as described above. Comprehensive protocols are available on protocols.io (dx.doi.org/10.17504/protocols.io.g3gbyjw, doi.org/10.17504/protocols.io.gysbxwe).

## Results and Discussion

### Most complete set of circadian clock orthologs in Cyanobacteria

Circadian rhythms are reported for many Cyanobacteria and first sequence analyses revealed that the core genes, known from *Synechococcus* 7942, are conserved in almost all cyanobacterial species [4, 53, 55, 83]. Even though daily rhythms seem to be conserved in Cyanobacteria, composition and quantity of corresponding genes on the genome level show high variability [54]. Dvornyk and colleagues first attempted to describe the variety of cyanobacterial core circadian clock systems in 2003 [53]. Since then, the amount and depth of sequencing data increased manifold, which allowed us to perform a detailed analysis of the KaiC-based circadian clock including the input and output pathways. The circadian clock proteins of *Synechococcus* 7942 (Table S1) and the three diverged KaiB and KaiC homologs from *Synechocystis* 6803 (Table S1) were the basis for our analysis. Their protein sequences served as basis for a reciprocal best hit BLAST analysis. Organisms with at least one homolog to KaiC were retained for further analysis. This stringent filter was essential since KaiC represents the core of the circadian clock in *Synechococcus* 7942. These organisms were grouped by their corresponding genus for a first overview of the homolog distribution (Fig. 1A). We found homologs in Cyanobacteria, Proteobacteria, Archaea, as well as other Bacteria such as *Chloroflexi*. This finding is in good agreement with previous studies [53, 55, 84]. However, our comprehensive study identified a plethora of new bacterial and archaeal genera harboring homologs to the circadian clock genes (Fig. 1A). Nevertheless, Cyanobacteria represent the phylum with the highest degree of sequence similarity and integrity of the system followed by Bacteria, mostly Proteobacteria. In Archaea, homologs to only a fraction of core genes could be identified (Fig. 1A).

**Figure 1:**
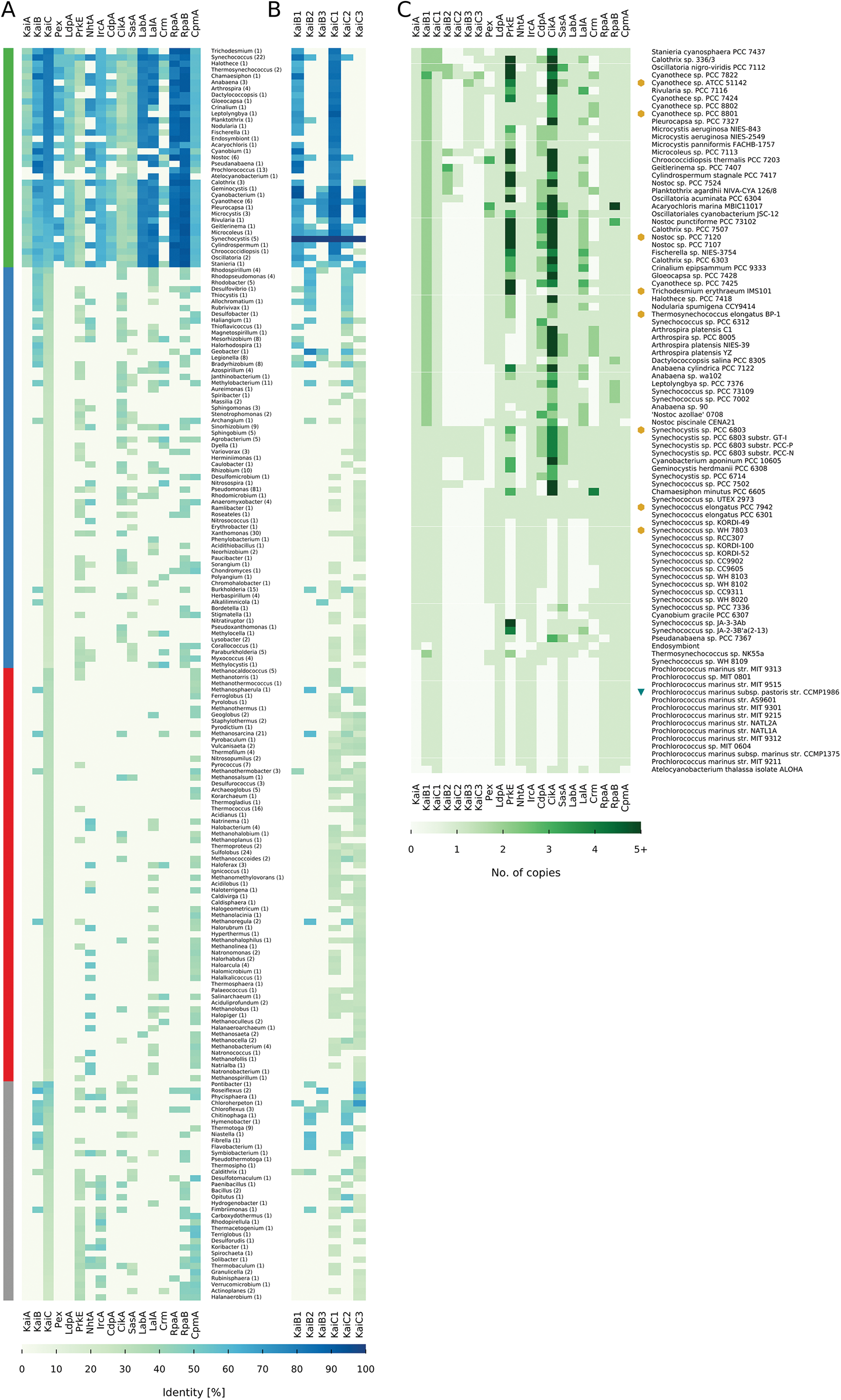
Distribution of circadian clock proteins. (**A,B**) Shown are the mean sequence similarites of each protein for each genus that contains a KaiC homolog from *Synechococcus* 7942. The genera are sorted by their group and KaiC similarity. The number in parenthesis represents the number of individual genomes per genus. (**A**) shows the mean similarities for the circadian clock proteins of *Synechococcus* 7942. (**B**) shows the mean similarities for the circadian clock proteins of *Synechocystis* 6803. The four taxonomic main groups are highlighted: Cyanobacteria (Green), Proteobacteria (Blue), Archaea (Red), Other (Grey). (**C**) shows the number of homologs for each protein in each cyanobacterium. Copy numbers higher than five were condensed. Cyanobacterial strains with true circadian clocks [87, 73, 74, 88, 75, 89, 90, 8, 91, 92, 93] are highlighted with a yellow hexagon and the strain with hourglass like timing mechanism [94] with a green triangle. We could not find references for all strains, however, we identified strains with circadian rhythms, which are not listed here as their genomes are still not fully assembled.

### Core circadian clock factors KaiB and KaiC beyond the phylum of Cyanobacteria

Four out of seventeen studied factors are exclusively found in Cyanobacteria (Fig. 1A). One of these factors is KaiA, as previously reported in studies with a smaller sample size [53, 84]. However, even some Cyanobacteria, like *Candidatus Atelocyanobacterium thalassa* (previously named Cyanobacterium UCYN-A), and all representatives from the genus *Prochlorococcus* lack *kaiA* (Fig. 1A) [54]. Interestingly, multiple copies of *kaiA* in a single cyanobacterial genome could not be identified. This is of special interest, because we could observe strong sequence length variations for KaiA (Fig. 5A, C) and multiple copies for the other core proteins KaiB and KaiC have been reported (Fig. 1B) [53, 54, 55]. With the *kaiA*-lacking *Candidatus Atelocyanobacterium thalassa* and a *kaiA*-containing cyanobacterial endosymbiont [85] only two out of 94 studied Cyanobacteria do not contain a *kaiB*, whereas a third of the cyanobacterial genera contain multiple copies of *kaiB* and *kaiC* (Fig 1B, C). There are a few exceptional cyanobacteria like *Gloeobacter violaceus* [86], which are even lacking the *kaiC* gene and are thus not detected in this analysis due to the previously described filtering criteria. However, all Cyanobacteria having homologs to *kaiB* and *kaiC* are coding for at least one pair of proteins most similar to KaiB1 and KaiC1 from *Synechocystis* 6803 (Fig. 1B, C), which is consistent with previous studies [53, 55]. Only *Cyanothece* sp. PCC 7822 has homologs similar to all three *kaiB* and *kaiC* copies from *Synechocystis* 6803 (Fig. 1C). The majority (67.68%) of the bacterial genera, outside of the cyanobacterial phylum, encode KaiC3-like proteins. Approximately half of them also contain additional KaiC homologs, most similar to KaiC1, KaiC2, or sometimes even both (Fig. 1B). This observation does not hold true for KaiB homologs, here KaiB2 is the major KaiB homolog (75.86%). One group of organisms, all belonging to the phylum of *Bacteroidetes*, stands out in this analysis, because they encode only proteins similar to KaiB2/KaiC2 (Fig. 1B). Archaeal genera show mainly homologs with highest identity to KaiC1 or KaiC3. Furthermore, almost all of the Archaea have multiple copies of the KaiC protein. Whereas only four have homologs to KaiB, which is either similar to KaiB2 or in thermophilic *Methanothermobacter* similar to KaiB1 (Fig. 1B).

### Circadian clock factors solely involved in cyanobacterial input pathways

Multiple clock input factors are present in all four taxonomic groups namely PrkE, NhtA, and CikA. The latter also acts in the output of the circadian clock (Table S2, Fig. 1A) [36, 37, 44]. The prevalence of CikA is in good agreement with its tremendous effect on resetting the circadian clock in *Synechococcus* 7942 [37], where its interaction with the Kai complex is mediated by additional proteins [36, 38]. CikA destabilizes in the presence of oxidized quinones and thereby integrates information about the cellular redox state into the oscillator [36]. Naturally, CikA-lacking *Prochlorococcus* are not able to reset their timing mechanism in continuous light [94], pointing at the importance of this factor. However, single mutations in *sasA* can restore circadian properties in *Synechococcus* 7942 *cikA* mutants and it has been suggested that a simpler network with modified interactions of the other clock proteins can exist [95]. The input factors Pex, LdpA, and CdpA are found to be unique for cyanobacteria (Fig. 1A), confirming a previous analysis of *ldpA* [96]. Hence, with Pex, a transcriptional repressor of *kaiA* [34], and LdpA, a redox-sensing protein [20] two factors, which sense the cellular metabolic state of the circadian clock in *Synechococcus* 7942, are missing outside of the cyanobacterial phylum. LdpA is also the only input factor present in the reduced but functional timing system of *Prochlorococcus* [54, 83]. This might indicate the necessity of this factor for the entrainment of the clock [60]. Nevertheless the possibility of a functional clock without Pex and LdpA remains, since LdpA and Pex mutants in *Synechococcus* 7942 are only altered in their period length [97, 35] and Pex is also missing in the KaiA-lacking *Prochlorococcus* and the KaiA-containing *Synechocystis* 6803 (Fig. 1A). In *Synechococcus* 7942 the third unique cyanobacterial input factor, CdpA, influences phase resetting and acts in parallel to CikA [38]. Since CdpA seems to be essential in *Synechococcus* 7942 [38], it likely has a prior role in processes other than phase resetting and is hence dispensable for input pathways in other organisms than Cyanobacteria.

Altogether, the absence of three important input factors outside of the cyanobacterial phylum suggests that other entrainment systems might be used for putative timing systems. In *Rhodobacter sphaeroides*, which displays circadian gene expression rhythms, a histidine kinase is encoded in an operon with *kaiBC* and was suggested as a candidate for transducing the redox signal to KaiBC [98]. Further, the direct entrainment by the ATP/ADP ratio [30, 99] might be the primary mechanism to synchronize the circadian clock with metabolism and the environment.

### Central output factors are missing in Archaea and non-cyanobacterial genera

The output pathway of the circadian clock in *Synechococcus* 7942 involves eight proteins (see Fig. 7). RpaA serves as a key regulator [43]. Its activity is indirectly modulated depending on the phosphorylation state and the ATPase activity of KaiC [44, 100]. SasA (antagonistically to CikA) connects the core clock to RpaA, which in turn regulates global gene expression, including the *kaiBC* promoter [41, 43, 44, 101]. LabA, CrmA and RpaB are also known to affect RpaA [42, 46, 47, 50, 101]. CpmA modulates *kaiA* expression by an unknown mechanism [49]. In contrast to the unique cyanobacterial input factors, we found none of the eight output proteins exclusively in Cyanobacteria. Homologs of five factors (SasA, CikA, LalA, Crm, CpmA) are present in all four investigated taxonomic groups (Table S2). CpmA is a member of a superfamily essential for purine biosynthesis and thus likely to have orthologs in other organisms [49]. However, RpaA and RpaB are not present in Archaea and SasA is only found in the methanogenic genera *Methanospirillum*, and *Methanosalsum*. Hence, the entire central output pathway is missing in Archaea (Fig. 1A). In addition, orthologs for RpaA are found in only nine non-cyanobacterial genera, questioning whether another transcription factor might read out the putative core timer in other Bacteria (Fig. 1A). A previous BLAST search by Dvornyk and colleagues revealed that SasA homologs in non-cyanobacterial prokaryotes lack the KaiB-like domain [61]. This finding is confirmed in our analysis, indicating that stimulations of SasA homologs by KaiC outside of Cyanobacteria are very unlikely, because interaction occurs via this KaiB-like domain, which adopts a thioredoxin-like fold [39, 102]. Altogether, our analysis reveals that possible circadian clocks of Bacteria and Archaea must use an output pathway that is different from the one described in *Synechococcus* 7942.

### Co-occurrence analysis hints at the core module for circadian timing

The previous analysis revealed substantial differences in the composition of the clock components between Cyanobacteria, other Bacteria, and Archaea. Even within Cyanobacteria there is a huge variety in the composition of the potential circadian clocks (Fig. 1C). Cyanobacteria have either a severely reduced timing systems, such as the one in *Prochlorococcus*, a standard system as seen in *Synechococcus* 7942, or an inflated system as found in *Synechocystis* 6803 (Fig. 1C). This trichotomy of systems raises questions about essentiality and pairwise co-occurrence of circadian clock proteins. These questions were answered in a series of right-sided Fisher’s exact tests. To avoid systematic biases due to an overrepresentation of closely related strains [103], we extracted 69 unique combinations of the 21 circadian clock factors as described in materials and methods.

Within these 69 unique systems two factors are always present: (i) KaiC, because we selected for organisms containing at least one KaiC-like protein and (ii) RpaB, which is associated with cell size and circadian gene expression [42, 51]. RpaB competes for promoter binding sites with RpaA and its phosphorylated state is thought to inhibit the phosphorylation of RpaA [47]. Other factors present in the majority (≥ 90 %) of the observed unique clock systems are KaiA, KaiB, LdpA, IrcA, CikA, SasA, RpaA and CpmA. Because of their abundance, most of these factors show no pairwise co-occurence. For example RpaA and RpaB are found in 68 and all 69 clock systems, respectively. Thus their joint presence comes to no surprise. Instead, the finding confirms essential roles in global transcription regulation of Cyanobacteria [104]. With Fisher’s exact test we seek to identify gene pairs rather unexpectedly co-occuring in a smaller subset of organisms. Such findings can indicate a common function in the circadian clock system. Only KaiA and CikA, out of the most abundant factors, show significant co-occurence with other factors.

Within the input pathway we detected three siginficantly co-occuring pairs (Fig. 2): (i) CikA and its interaction partner PrkE [38] (ii) PrkE and CdpA and (iii) the *kaiA* repressor Pex and CdpA. The first two results are in good agreement with a previous study [38]. Within the output pathway, there is a significant co-occurrence between LabA and its ortholog LalA. Interestingly, we identified several significant co-occurrences between factors of the input and the output pathway. CikA, which functions in the input and output of the clock, co-occurs significantly with LabA, and LalA (Fig. 2). This fits well in the overall picture as CikA and LabA are thought to regulate the activity of RpaA [44, 50]. Additionally, PrkE shows also significant occurrences with both LabA and LalA (Fig. 2). Furthermore, CdpA was found to co-occur significantly with LabA. (Fig. 2) This is of special interest since PrkE and CdpA are only known as interaction partners of CikA, and both are involved in phase resetting, and cell division, respectively [38]. This result, however, hints at potential increased involvement of PrkE and CdpA in the RpaA regulation and supports the view of an integrated network with overlapping interactions of input and output factors [105]. Notably, no co-occurrence of NhtA and LdpA was detected, although it was suggested that NhtA might be involved in assembly of the iron-sulfur cofactor of LdpA [38]. Significant co-occurrences with core factors could only be observed between KaiA and LalA (Fig. 2). However, KaiA shows a strong, but not significant, co-occurrence with CikA (p = 0.0105). Lastly, we also found significant co-occurrences between KaiB2 and KaiC2 as well as KaiB3 and KaiC3 (Fig. 2). This indicates two distinct function of the two pairs. In this context it is worth mentioning that *kaiB2*-containing archaeal genomes always encode a KaiC2 homolog.

**Figure 2:**
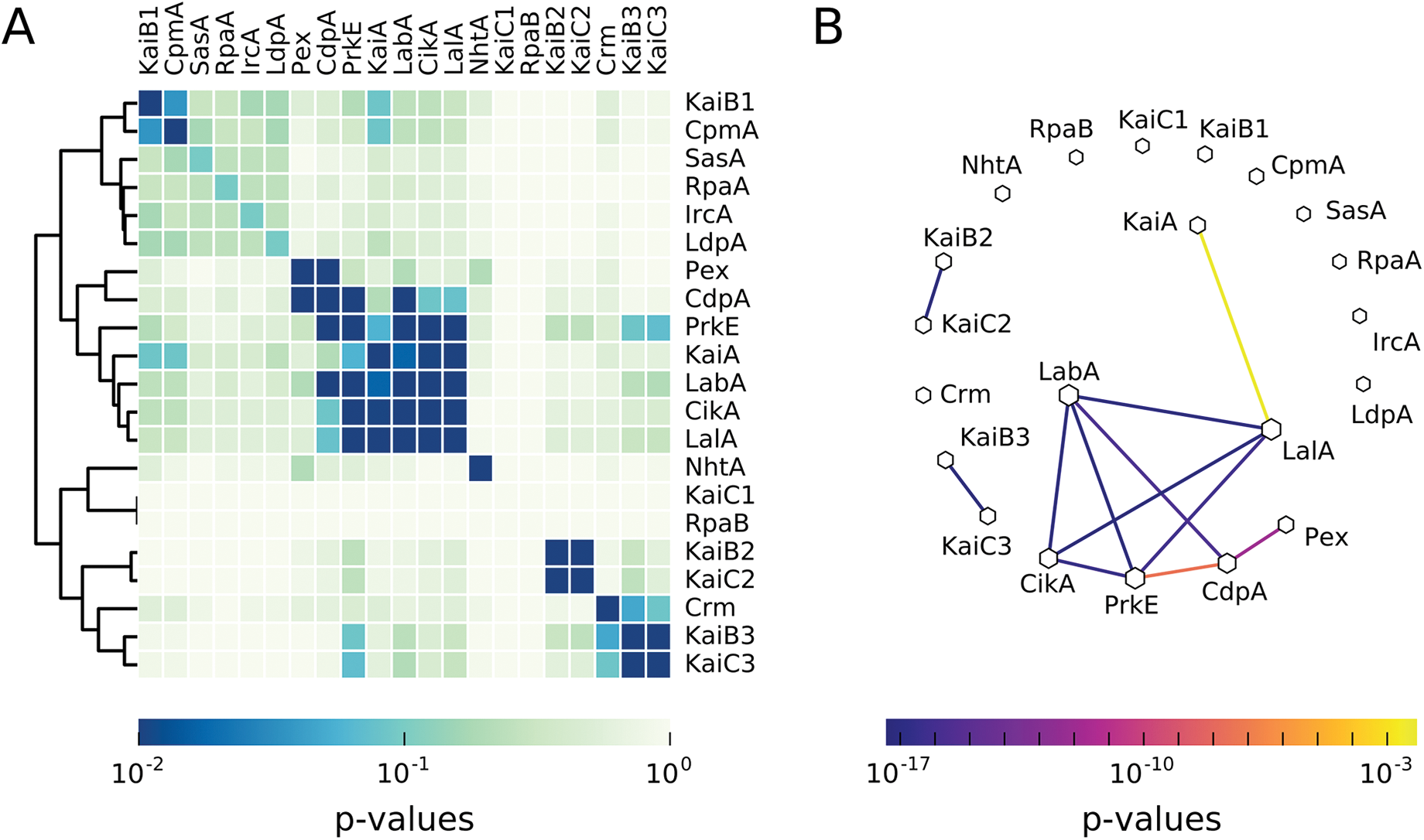
Co-occurrence of circadian clock proteins in cyanobacteria. (**A**)The p-values, calculated by pair-wise Fisher’s exact tests, are visualized in a heatmap. Only p-values ≤ 0.01 are considered as significant. Proteins are sorted by a hierarchical agglomerative clustering algorithm. (**B**) Network of significant co-occurring circadian clock factors in cyanobacteria, calculated in regard to the results of the pair-wise Fisher’s exact test. The line color corresponds to the level of significance. Missing links are those that had a higher p-value than 0.01. Node size is proportional to the degree of that node.

In summary, we identified a conserved set of factors (Fig. 2), both in input and output that show significant co-occurrences. This set, composed of KaiA, PrkE, CdpA, CikA, LabA, and LalA, is found in Cyanobacteria with a true circadian clock such as *Synechococcus* 7942 and *Synechocystis* 6803 [73] but is missing in cyanobacterial strains with reduced timing mechanisms such as *Prochlorococcus*. Additionally, PrkE, CikA, LabA, and LalA are also missing in most marine *Synechococcus* species. This finding hints at the importance of these factors for the functionality of a circadian clock. On the other hand, NhtA and Crm seem to play only a minor or extending role in clock regulation, because they are neither always present nor show significant co-occurrence with other factors.

### A systematic analysis of circadian expression in Cyanobacteria

Genome-wide time-resolved expression measurements in a range of cyanobacterial strains have repeatedly indicated substantial fractions of genes with circadian regulation patterns [106, 107, 87]. Considering that all Cyanobacteria share the challenge of a photoautotrophic lifestyle, which requires major changes in the metabolism between day and night, one might expect a common transcriptional regulatory pattern. Thus, we compared a total of nine published microarray time-series datasets of different cyanobacterial strains under constant light or diurnal light conditions (for details see '
Table S3), which were available and applicable for this analysis. Not all of the chosen microarray experiments were conducted under constant light conditions, which leads to a combination of circadian-clock regulated and light-induced genes. We therefore refer to genes with oscillating expression as diurnally regulated instead of circadian. For allowing a direct comparison, we reprocessed the raw-data and subjected the resulting expression time series to a harmonic regression oscillation detection. This method assumes a sinusoidal shape of circadian expression profiles and uses linear expression profiles as background, yielding estimates of the peak phase and amplitude of each gene.

In a first step we compared biological replicate datasets to establish the reproducibility of strain-specific circadian expression programs. Similarity between two circadian expression programs was established using the circular correlation coefficient *ρ*ccc as described by Jammalamadaka and Sarma [108] applied to estimated peak expression phases. The following analyses were limited to genes with oscillating expression profiles in both compared datasets since only in these cases phase and amplitude estimates are meaningful descriptors. Direct comparison of the oscillation phases and amplitudes indicates good reproducibility between two respective measurements in form of statistically significant elevated correlation of the diurnal expression patterns in *Synechococcus* 7942 (*ρ*ccc = 0.61, p ≪ 0.01), *Synechocystis* 6803 (*ρ*ccc = 0.31, p ≪. 0.01), and *Cyanothece* sp. ATCC 51142 (*ρ*ccc = -0.51, p ≪ 0.01) (Figure 4 top row). While the *Synechocystis* 6803 datasets show significant similarity, the correlation is diminished by the distinct concentration of expression phases during the day in the beck14 [109] dataset compared to the leh13 [76] measurements. Interestingly, both *Cyanothece* sp. ATCC 51142 datasets exhibit a good agreement of peak expression phases with large early day and early night clusters, but the large negative correlation emphasizes the presence of a significant number of anti-phasic gene pairs. The corresponding oscillation amplitude values exhibit high statistically significant correlations (*ρ*ccc *>* 0.77) for all three datasets (Fig. 3 bottom row).

**Figure 3:**
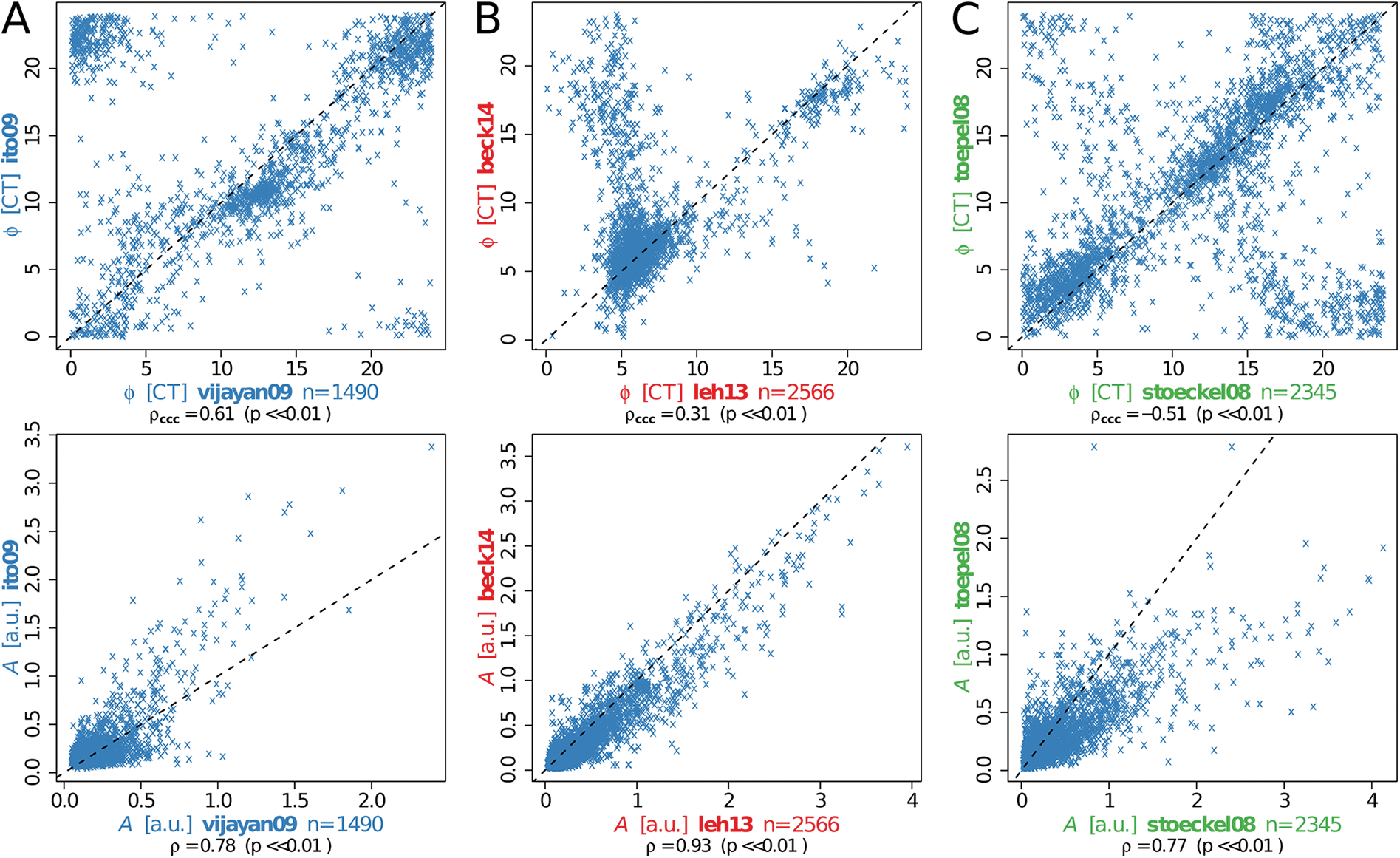
Phase and amplitude reproducibility of diurnal genes within cyanobacterial strains. Comparison of expression phase *φ* (top, [CT]) and amplitude A (bottom, [a.u.]) of diurnal genes shared between independent datasets of the same cyanobacterial strain. (**A**) *Synechococcus* datasets of Vijayan and colleagues [119] (x-axis) and Ito and coworkers [87] (y-axis), (**B**) *Synechocystis* in the datasets of Lehmann and coworkers [76] (x-axis) and Beck and colleagues [109] (y-axis), and (**C**) *Cyanothece* in St¨ockel and colleagues [120] (x-axis) and Toepel and colleagues [121] (y-axis). The number of genes n found to oscillate significantly and the corresponding Pearson correlation coefficient *ρ* between *φ* and A is provided below each panel, followed by the respective p-value for *ρ* differing from 0. For *φ*, the circular correlation coefficient *ρ*ccc is also provided. Axis labels are shown in the strain-specific color.

This observation motivated the second step of the analysis, the comparison of expression patterns between different cyanobacterial strains. To facilitate this comparison, the prediction of homologous genes in the cyanobacterial clade by Beck and colleagues [109] was used as starting point. We focused first on the set of genes with oscillating expression patterns in all datasets. The estimate of the core diurnal genome based on the presented data collection spans 95 genes (Table S5), which are mostly involved in central metabolic processes. Most strikingly, 18 out of 64 genes annotated by Cyanobase [68] as “ribosomal protein” genes in *Synechocystis* 6803 fall into the core diurnal set, furthermore seven out of 27 genes annotated with “photosystem II”. The remaining diurnally expressed genes are found interspersed across the metabolic network. While genes coding for parts of photosystem II, the RNA polymerase, or the ribosomal proteins can be expected to serve important roles in the adaptation to photic and aphotic phases, this analysis ascribes similar importance to other metabolic processes e.g. in the repair of UV-damaged Photosystem II centers (*slr1390*) [110], in the phosphate transport system (*pstB2*), pyrimidine and arginine biosynthesis (*sll1498*), or the glycolysis/gluconeogenesis via the fructose-bisphosphate aldolase (*sll0018*). Marker gene candidates for a working clock can be derived from the core diurnal genome. Importantly, less confidence can be placed in genes which exhibit peak expression phases shortly past dawn since these can either be clock regulated or simply induced by light. Two high confidence candidates are the light-independent protochlorophyllide reductase subunit ChlB and the fructose-bisphosphate aldolase *fbp*. Indeed, *fbp* also shows circadian expression patterns in *Clamydomonas reinhardtii* [111] and *Arabidopsis thaliana* where its late-night peaks may reflect the great importance of these aldolases in higher plants for the mobilization of plastidic starch [112]. Particularly in higher plants, the mobilization of starch, the conversion into sucrose, and its transport to other parts of the plant occur mainly at night.

The group of 15 “hypothetical protein” genes in the core diurnal genome constitutes an excellent candidate set for novel clock-driven genes in strains with a working core clock. Interestingly, several of these genes are implicated with cell division, such as the YlmG-related hypothetical gene *ssl0353*, which is required for proper distribution of nucleoids in cyanobacteria and chloroplasts [113]. Similarly, the hypothetical protein *slr1577* is suggested to function in the separation of chromosomes during cell division (Uniprot entry P74610). For the gene *slr1847* (Uniprot entry P73057) a DNA binding capability is suggested, which could therefore regulate expression, aid nucleoid organization, or protect the DNA.

The core clock genes *kaiA*, *kaiB*, and *kaiC* are a good starting point for a detailed comparative expression analysis. Only the *kaiB1* is significantly oscillating in all considered datasets. Interestingly, the *kaiA* gene features only very low amplitude expression oscillations and is arrhythmic in the vijayan09 *Synechococcus* 7942 dataset. The expression phases vary from dawn (*Microcystis aeruginosa* PCC 7806) to morning (*Synechocystis* 6803), over midday (stoeckel08 dataset of *Cyanothece* sp. ATCC 51142), and dusk (ito09 dataset of *Synechococcus* 7942), into night (*Anabaena* sp. PCC 7120). The observed expression phases of *kaiB1* are comparable to those of *kaiA*, but with significantly larger amplitude in *Synechococcus* 7942 datasets. The phase of the *kaiB1* homolog in *Prochlorococcus marinus* MED4 peaks before dawn, comparable to *Anabaena* sp. PCC 7120. The *kaiC1* expression phases and amplitudes match those of *kaiB1*, with the notable exception of *Cyanothece* sp. ATCC 51142 for which antiphasic late-night peaks are observed. In *Prochlorococcus marinus* MED4, *kaiC1* peaks during the early night in contrast to the late night phase of *kaiB1*.

Many aspects agree well with previous knowledge. In *Synechococcus* 7942, the core clock genes *kaiB* and *kaiC* are arranged in the *kaiBC* operon resulting in similar expression patterns [114, 115, 116], while *Cyanothece* sp. ATCC 51142 features the *kaiAB1C1* operon [117]. Interestingly, *Cyanothece* sp. ATCC 51142 features consistent anti-phasic expression of *kaiB1* and *kaiC1* whereas the remaining strains show co-expression, hinting at *Cyanothece*-specific post-transcriptional regulation of *kaiB1* or *kaiC1*. Oscillations in the *kaiA* gene expression, as reported by Ishiura and colleagues, feature small expression amplitudes compared to *kaiB* and *kaiC* [118]. In fact, *kaiA* consistently falls below the threshold of 2-fold expression change for the classification as circadian oscillator, which is commonly employed in microarray studies.

In the second step we generalized the detailed analysis of expression phases as presented for the core diurnal genome. We applied the circular correlation measure to all possible combinations of expression datasets. The resulting distribution reveals a clear separation between pairs of biological replicate datasets, featuring large numbers of shared oscillating genes and more extreme correlations (Fig. 4 red), and pairs of different cyanobacterial strains with fewer shared oscillating genes and much less extreme correlation coefficients (Fig. 4 blue). The only exception to this separation is the *Cyanothece* sp. ATCC 51142 dataset stoeckel08 (Fig. 4 green), which shares many oscillating genes with both *Synechocystis* 6803 datasets (leh13, beck14). The corresponding correlation coefficients are, however, similarly small compared to other inter-strain pairs. The full set of pairwise phase comparisons, which underlay this analysis are shown in Figure S1. This result indicates that the diurnal peak expression phase is not preserved amongst homologous genes in the cyanobacterial clade but might instead be tuned according to the metabolic gene outfit and the environmental needs of the respective strain.

**Figure 4:**
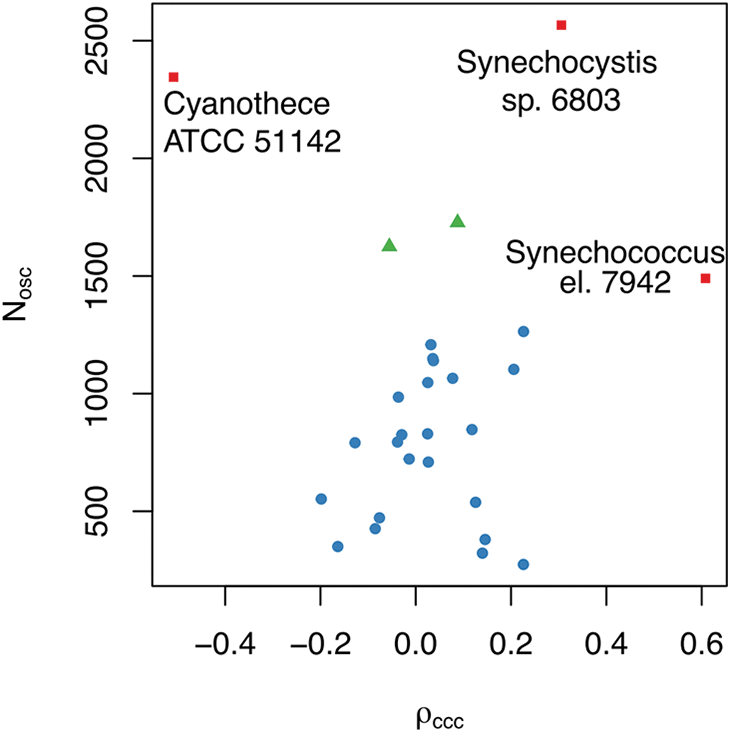
Expression phase similarity versus number of shared oscillating genes. The comparison is carried out between all dataset combinations. Biological replicate experiments available for three cyanobacterial strains show a large number of shared oscillating genes as well as large correlation coefficients (red). The comparison of the stoeckel08 dataset (*Cyanothece*) with both *Synechocystis* datasets is marked in green. The remaining comparisons between various strains are shown in blue.

### Core circadian clock proteins, KaiA, KaiB, KaiC, vary in number and length

The previously observed diversity of circadian clocks within Cyanobacteria, and between other Bacteria and Archaea prompted further sequence analyses of the core clock proteins KaiA, KaiB and KaiC. Length comparisons gave rise to some new features of variations between the core factors of the circadian clock (Fig. 5). As described in the preceding, our BLAST analysis detected KaiA exclusively in Cyanobacteria. Interestingly, we could distinguish three subtypes of KaiA. While the sequence length of most KaiA is around 300 amino acids (AA) (*Synechococcus* 7942: 284 AA) some stains have shortend homologs with a length of roughly 200 and 100 AA, respectively (Fig. 5A, C). Truncated KaiA proteins are almost exclusively found in members of the order *Nostocales*. Similar results were reported previously by Dvornyk and colleagues, who also observed a higher degree of polymorphism for the *kaiA* gene in comparison to *kaiB* and *kaiC* [52, 53]. Multiple alignments of the KaiA proteins (Fig. S2) verified that the truncated KaiA proteins have a shortened N-terminal sequence, which functions in the complete protein as the amplitude amplifier [122]. However, all of these KaiA orthologs contain the C-terminal part important for clock oscillation [122].

**Figure 5:**
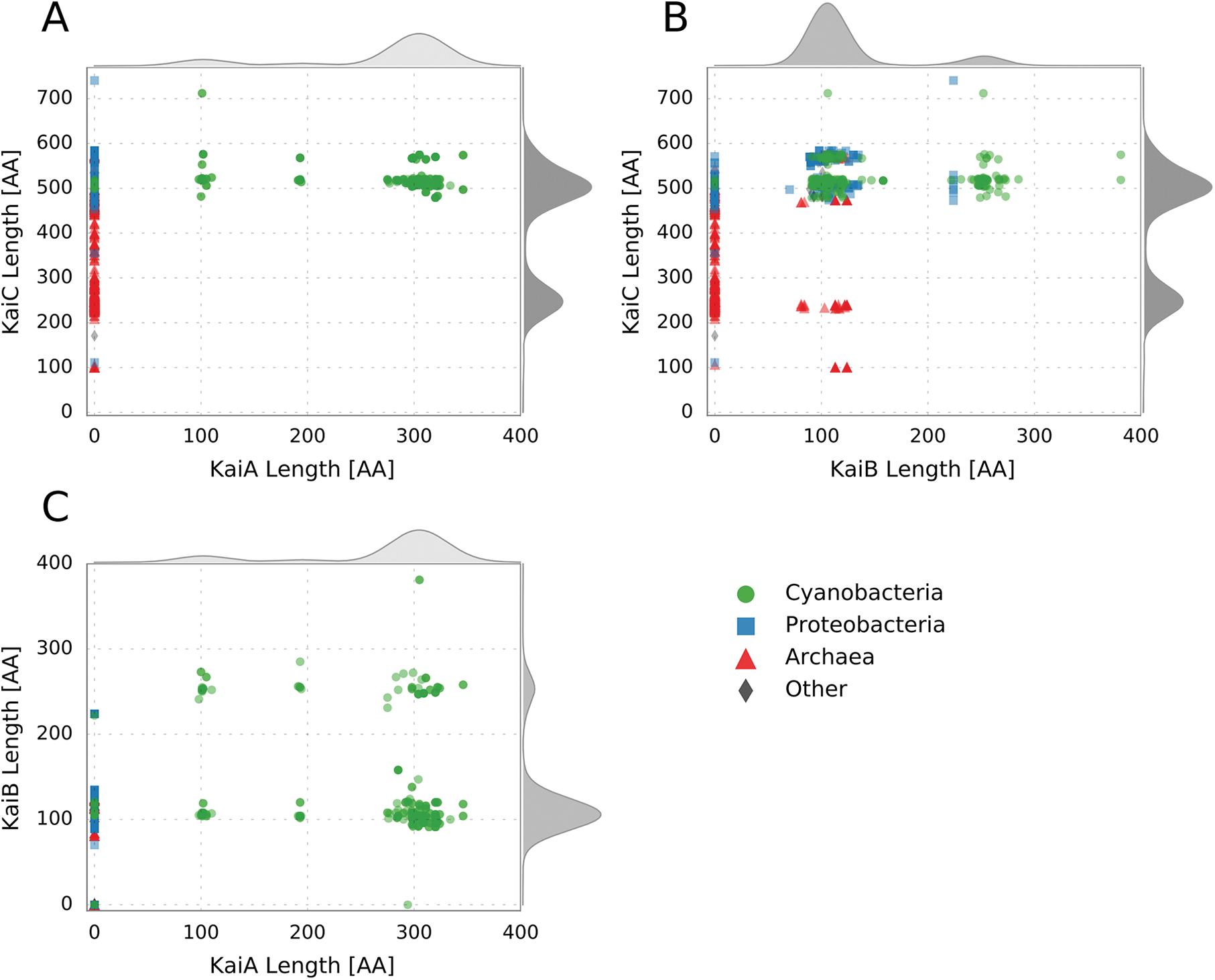
Protein length distribution of the circadian clock factors from *Synechococcus* 7942 and its homologs. The four taxonomic main groups are highlighted: Cyanobacteria (Green), Proteobacteria (Blue), Archaea (Red), Other (Grey). The curves outside of the plot represent the cumulative density distribution of the respective protein. (**A**) KaiC length distribution in dependency of the KaiA length. (**B**) KaiC length distribution in dependency of the KaiB length. (**C**) KaiB length distribution in dependency of the KaiA length.

Distribution of the KaiB protein length reveals two distinct groups. The KaiB homologs are either as long as the one from *Synechococcus* 7942 (102 AA) or about 250 AA in length (Fig. 5B, C). *Microcoleus* sp. PCC 7113 even has a KaiB with the length of 381 AA. KaiB homologs with query length are present in all four groups (Fig. 5B). Elongated KaiB proteins are mainly present in Cyanobacteria in the subclass *Oscillatoriophycideae* and the order *Nostocales*, specifying findings of Dvornyk [53] (Fig. 5C). BLAST analyses using the KaiB homologs from *Synechocystis* 6803 revealed that elongated variants are most similar to KaiB1. Elongation via concatenation of two KaiB was ruled out by visually inspecting alignments with two artificially concatenated KaiB1. Instead the elongated KaiB1 have a ~150 AA N-terminal extension. BLAST searches of the N-terminal region showed no homologous sequences in other organisms than Cyanobacteria, and no putative conserved domains could be identified. However, the N-terminal part is highly conserved within the Cyanobacteria having this KaiB variant. Interestingly, those *Nostocales* with an elongated KaiB1, also show a truncated KaiA.

The KaiC protein from *Synechococcus* 7942 is 519 AA in length and is build up by two domains, the CI and CII domain, which have a high similarity and are connected by a linker-domain [123, 124]. The C-terminal CII domain of KaiC comprises the interaction sites with KaiA as well as the specific phosphorylation sites [29, 125, 126, 127]. KaiC homologs were detected with lengths varying between 101 AA and 741 AA (Fig. 5A, B). There are KaiC homologs in Archaea representing the whole observed length spectrum of KaiC, whereas bacterial KaiCs are almost always about 500 AA in length (Fig. 5A, B). Furthermore, in Bacteria and Archaea KaiB (and KaiA in Cyanobacteria) is only found when a ‘full-length’ KaiC is present. In these bacterial organisms the length of KaiC is almost constant, regardless of the length of KaiA and KaiB homologs (Fig. 5A, B). However, in KaiB-possessing Archaea additional shorter KaiC homologs are found (Fig. 5B).

Moreover, the length distribution of KaiC revealed a substantial amount of KaiC homologs with a length of circa 250 AA, which is approximately the length of one KaiC domain. This KaiC variant is mainly found in Archaea, but also in a few Bacteria. In these bacterial species no KaiB homolog could be identified. Shorter KaiC homologs do not contain the important phosphorylation sites for maintaining the oscillator function. Therefore, they might not restore the full functionality of the *Synechococcus* 7942 KaiC, but can rather answer questions about the evolution of KaiC [53]. Regarding the evolution of KaiC, two valid hypotheses exist, both of which state that KaiC arose from a shorter ancestral *recA* gene followed by a gene duplication and fusion. However, on the one hand Leipe and colleagues [128] hypothesize that an ancestral single-domain KaiC originated in Bacteria, was transferred into Archaea, where its two-domain version originated, and a second lateral transfer event introduced the double domain KaiC into cyanobacteria. On the other hand Dvornyk and coworkers [53] argue in a follow up study that KaiC has to be of cyanobacterial origin. Given the amount of new genomic data further studies would help to unravel the evolutionary history of KaiC.

### Conserved motifs and activities in the cyanobacterial KaiC subgroups

For KaiC2 homologs outside of the cyanobacterial phylum an involvement in stress response (*Legionella pneumophila* [129]) and adaptive growth under rhythmic conditions (*Rhodopseudomonas palustris* [57]) has been demonstrated. Both proteins display autophosphorylation and KaiC2 from *Rhodopseudomonas palustris* shows elevated ATPase activity [57]. Nevertheless, the function of cyanobacterial KaiC2 and KaiC3 homologs remains unclear. We already demonstrated that KaiC2 and KaiC3 from *Synechocystis* 6803 displays kinase activity, which is independent of KaiA, whereas KaiC1 behaved like its *Synechococcus* 7942 ortholog [55]. Those activities could also be predicted from the C-terminal amino acid sequences [55]. To test whether general features of the three KaiC subgroups can be predicted, multiple alignments of the cyanobacterial KaiC1, KaiC2 and KaiC3 sequences were constructed. A WebLogo analysis revealed that relevant motifs for phosphorylation and dephosphorylation in the CII domain are highly conserved. The ATP-binding Walker Motif A (P-loop in Fig. 6A, GXXXXGKT, [118, 130, 131]) is present in all three KaiC subgroups. Strikingly, the respective sequence of KaiC-7942 (GATGTGKT) shows almost no modifications in KaiC1 and KaiC3 proteins. Furthermore, catalytic glutamates (EE in Fig. 6A, [132, 133]), the R-finger contacting the *γ*-phosphate of ATP [134], and the truncated Walker motif B (WalkerB in Fig. 6A, [26, 118, 131]), were found in all cyanobacterial KaiC subgroups. Notably, the arginine residue of the *Synechococcus* 7942 Walker B motif is not conserved in KaiC2 homologs. Serine and subsequent threonine are the dominant phosphorylation sites in KaiC1 and KaiC3 proteins, like S431 and T432 in KaiC-7942 [125, 126], whereas KaiC2 homologs display two serine residues. In some KaiC3 homologs a tyrosin is present as second phosphorylation site. T426, which is important for dephosphorylation of KaiC-7942 [125, 132, 135, 136], is also highly conserved. Therefore, phosphorylation and dephosphorylation via autokinase [26], ATP synthase and ATPase activity [27, 132], respectively, are very likely for all cyanobacterial KaiC homologs. The same holds true for the N-terminal ATPase activity: We observed high conservation of the Walker motif A (P-loop in Fig. 6A), the catalytic glutamate residues (EE in Fig. 6A) and the R-finger in the CI domains of all cyanobacterial KaiC subgroups. The presence of the R-linker in CI domains of KaiC1 and KaiC3 homologs indicate a structural coupling of the N-terminal CI and the C-terminal CII-domain as it was demonstrated for *Thermosynechococcus* KaiC [134], whereas KaiC2 homologs lack the R-linker in CI.

**Figure 6:**
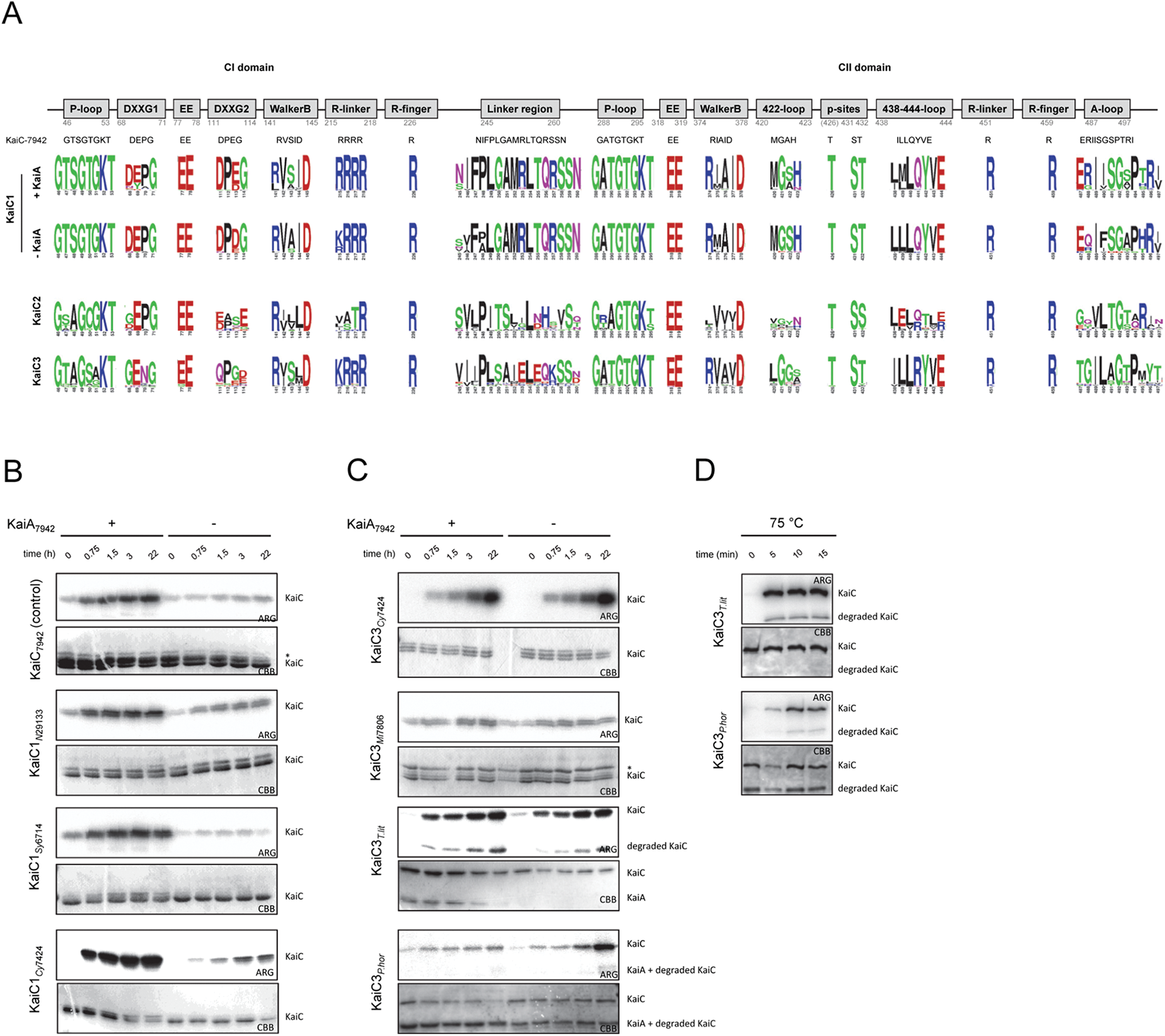
Activity of diverged KaiC homologs. (**A**) Conservation of important motifs in cyanobacterial KaiC1, KaiC2 and KaiC3 homologs based on a WebLogo analyses. Motifs in KaiC1 homologs are displayed for proteins from organisms, which encode a KaiA protein or lack KaiA, respectively. Numbers indicate the residues in KaiC-7942. Properties of the residues are displayed as follows: polar (green), neutral (purple), basic (blue), acidic (red), and hydrophobic (black). (**B**,**C**) Phosphate uptake analyses of selected KaiC1 (**B**) and KaiC3 (**C**) homologs at 30 *^◦^*C in dependence of KaiA. KaiC proteins were incubated with or without KaiA-7942 in the presence of *γ*-P^32^-ATP. After 0, 0.75, 1.5, 3, and 22 hours samples were separated via SDS-PAGE, stained with Coomassie (CBB), and subjected to autoradiography (ARG). The asterix indicates a contaminating protein. (**D**) Kinase activity of archaeal KaiC3 homologs at 75 *^◦^*C. KaiC3 proteins from *Thermococcus litoralis* and *Pyrococcus horikoshii* were incubated with *γ*-P^32^-ATP and autophosphorylation was analyzed after 0, 5, 10, and 15 minutes. Shown are the Coomassie stained proteins (CBB) and autoradiography (ARG).

KaiC homologs from the genus *Prochlorococcus* were classified as KaiC1 orthologs in our BLAST analysis (Fig. 1B). However, *Prochlorococcus* strains do not contain KaiA, and KaiC from *Prochlorococcus marinus* MED4 was demonstrated to phosphorylate independently of KaiA due to a modified A-loop sequence [83]. Therefore, we compared WebLogos of the A-loop sequence [28] for KaiC1 orthologs from Cyanobacteria with and without KaiA (Fig. 6A). The most obvious difference to KaiC1 proteins from cyanobacterial strains with KaiA is the presence of neutral glutamine in the second position, instead of a positively charged arginine. In KaiC2 and KaiC3 Weblogos motifs are even less conserved as already described for the *Synechocystis* 6803 representatives [55]. Interestingly, the KaiC3 WebLogo motif does not display any charged residue anymore. The absence of A-loop residues that are important to keep it in the buried state [28] indicate that the KaiA-independent phosphorylation is characteristic for KaiC2 and KaiC3 homologs. This is supported by the low conservation of the 438-444-loop and/or the 422-loop, which are part of the interaction network that mediates inhibition of phosphorylation by the buried A-loops in *Synechococcus* 7942 [28, 124]. The high conservation of these loops in KaiA-lacking strains remains enigmatic.

To test this hypothesis, phosphate uptake as an exemplary KaiC activity was analyzed for representative cyanobacterial KaiC1 and KaiC3 proteins by incubation with *γ*-P^32^-ATP at 30 *^◦^*C in the presence and absence of *Synechococcus* 7942 KaiA (KaiA-7942) The well-studied KaiC from *Synechococcus* 7942 (KaiC-7942) served as control. As demonstrated in Figure 6B and 6C all recombinant KaiC proteins incorporated phosphate over time. The intrinsic kinase activity of KaiC1 homologs from *Nostoc punctiforme* ATCC 29413 (KaiC1-N294133), *Synechocystis* sp. PCC 6714 (KaiC1-Sy6714), and *Cyanothece* sp. PCC 7424 (KaiC1-Cy7424) was stimulated by KaiA-7942, similar to KaiC-7942 (Fig. 6B). As expected, KaiA had no effect on autophosphorylation of KaiC3 from *Cyanothece* sp. PCC 7424 (KaiC3-Cy7424) and *Microcystis aeruginosa* PCC 7806 (KaiC3-Mic7806, Fig. 6C). To extend the analysis to non-cyanobacterial proteins, KaiC3 from the hyperthermophilic Archaea *Thermococcus litoralis* (KaiC3-T.lit) and *Pyrococcus horikoshii* (KaiC3-P.hor), which show optimal growth at 85 *^◦^*C and 98 *^◦^*C [137, 138], were analyzed in a similar way. Again both recombinant KaiC3 proteins displayed phosphorylation at 30 *^◦^*C, which was independent of KaiA-7942 (Fig. 6C). Incubation at 75 *^◦^*C indicated that the two archaeal KaiC3 homologs display kinase activity also at high growth temperatures (Fig. 6D). Hence kinase activity of KaiC proteins seems to be well-conserved, independent of the growth conditions of the strains, they are originating from.

## Conclusion

### A core module for circadian regulation

Our analysis of 11,264 genomes clearly demonstrates that components of the *Synechococcus* 7942 circadian clock are present in various bacteria and archaea. However, the frequency of Kai-clock related proteins is highest in Cyanobacteria. In fact KaiA, Pex, LdpA, and CdpA are exclusive to organisms of this phylum. In other organisms, e.g *Rhodobacter sphaerodies*, reduced KaiBC-based clock systems are likely able to drive circadian oscillations [98]. An even simpler system solely dependent on KaiC might enable diurnal rhythms in *Haloferax volcanii* [59], probably using the ATP/ADP ratio for clock entrainment. Predictions for KaiC activities based on sequence alignments and motif analyses were validated through biochemical experiments. We confirmed kinase activity for ‘full-length’ KaiC proteins composed of one CI and one CII domain, even in organism without *kaiA* or *kaiB*. KaiA from *Synechococcus* 7942 enhanced KaiC-phosphorylation only in strains naturally possessing a *kaiA* gene.

Our co-occurrence analysis hints to a conserved extension set for circadian regulation, which is present in cyanobacteria with observed circadian behavior and absent in cyanobacteria having a diurnal, hourglass-like lifestyle only (see also Fig. 7). A diurnal core set, which is important to enable an hourglass-like timing system that resets every day, might be composed of KaiB, KaiC, LdpA, IrcA, SasA, RpaA, RpaB, and CpmA. However, our identified circadian core set, which potentially enables a selfsustained clock, additionally consists of KaiA, the two input factors CdpA, and PrkE as well as the input and output factor CikA, and the output factors LabA, and LalA.

**Figure 7:**
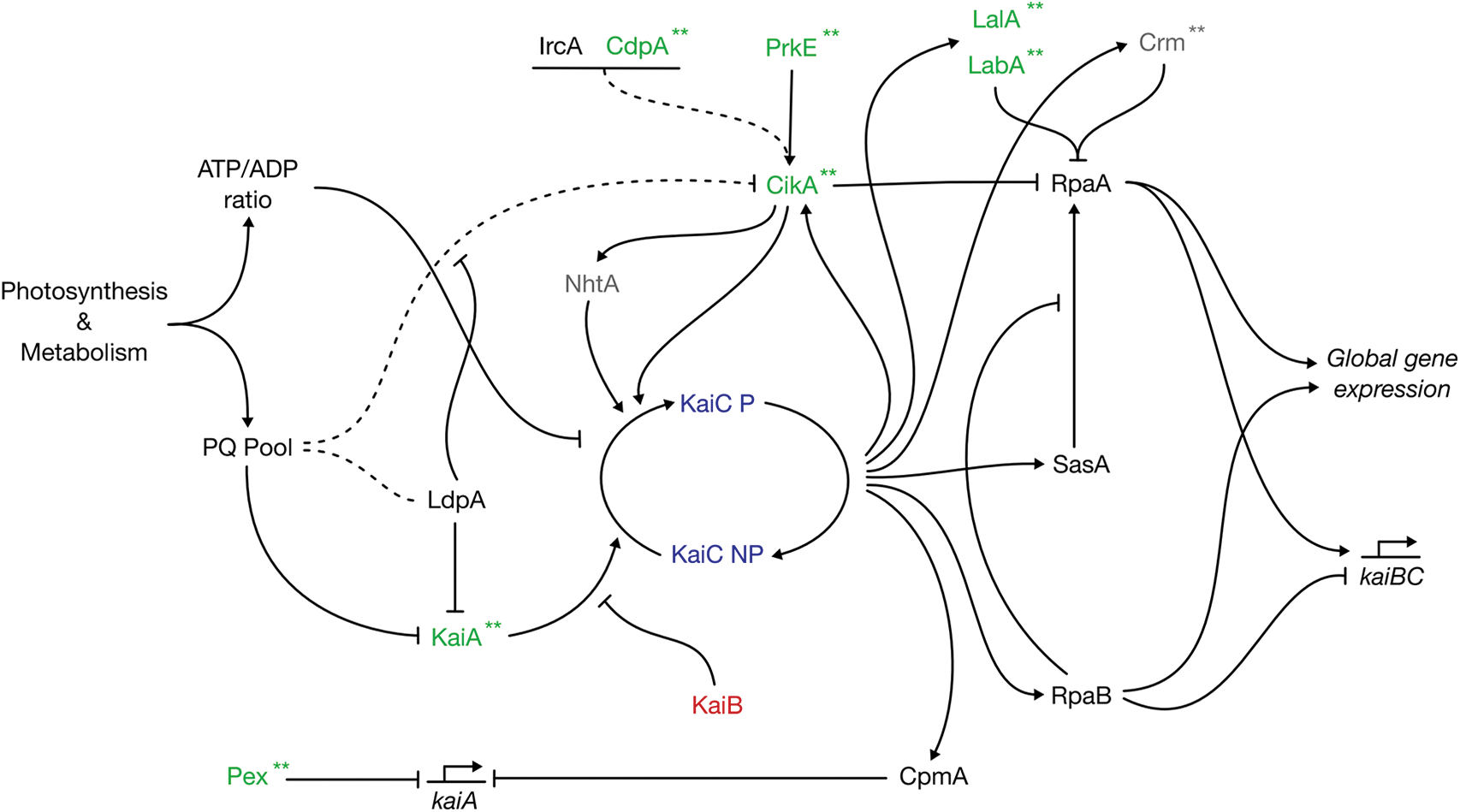
Schematic overview of the circadian clock in *Synechococcus* 7942. Shown are the protein interactions based on [20, 34, 35, 36, 37, 38, 39, 40, 41, 42, 43, 44, 45, 46, 47, 48, 49, 50, 51]. The plastoquinone pool and ATP/ADP ratio serve as input signals to entrain the circadian clock with the environment and metabolic state of the cell. The signals are recognized by the input factors LdpA and CikA and the core clock factors KaiA and KaiC. Other input factors are Pex, and the CikA interaction partner NhtA, PrkE, IrcA, and CdpA. The stimulating effect of KaiA on KaiC is antagonized by KaiB. The output of the clock is comprised by SasA, CikA, RpaA, RpaB, LabA, LalA, Crm, and CpmA. SasA interacts with KaiC and further phosphorylates RpaA, whereas CikA acts as a phosphatase on RpaA. RpaA is also regulated by LabA, Crm, and RpaB. RpaA together with RpaB controls global gene expression as well as the expression of the *kaiBC* cluster. CpmA is a transcriptional regulator of *kaiA*. The seven most interconnected factors we found in our co-occurrence analysis are highlighted in green. Factors colored in light grey showed no co-occurrence in the Fisher’s exact test and are present in *<* 90% of all observed systems. Asterisks (**) indicate the factors missing in *Prochlorococcus*.

The systematic comparison of microarray timeseries datasets indicates that the diurnal peak expression phase is not conserved amongst homologous genes in the cyanobacterial clade. Instead, the expression phase may be tuned according to the gene outfit and varying environmental needs. The analysis yielded a set of 95 genes in the core diurnal genome, which can be considered critical for the adaptation to day and night. Particularly the subset of non-light induced genes are prime candidates for circadian clock marker genes. This set furthermore contains several hypothetical genes, which are interesting candidates for novel clock-driven genes.

The gained insights about the diversity within the composition of the components involved in the circadian protein clock as well as the diversity on the sequence level of the core factors call for further modification and simplification of the clock. The exponential increase of molecular tools for synthetic applications in recent years sets the stage for such ambitious projects. A future goal could be the reduction of complexity by removing as many factors as possible so that an integration of a circadian clock in synthetic and industrially valuable organisms becomes feasible.

## Declarations

## Acknowledgement

We thank Mariam Yazdanyar for help with protein expression and purification. We are very grateful to Elke Dittmann, Wolfgang Lockau, and Wolfgang R. Hess for providing genomic DNA of *Microcystis aeruginosa* PCC 7806 and *Nostoc punctiforme* ATCC 294133, *Cyanothece* PCC 7424, and *Synechocystis* PCC 6714, respectively. This work was supported by DFG grants AX 84/1-3 and EXC 1028 to NMS, AW and IMA. ERC starting grant (Nr. 311523, Archaellum) for PC and SVA. RL was supported by the grant “CyanoGrowth” funded by the German Federal Ministry of Education and Research (reference: FKZ 0316192 to Ralf Steuer).

## Author contributions

IMA and AW supervised the study. NMS, CB, and RL designed and NMS and RL carried out bioinformatic data analyses. AW designed and performed sequence analysis and *in vitro* experiments. AW and PC cloned, expressed, and purified recombinant proteins. NMS, RL, CB, AW, and IMA prepared the manuscript. PC and SVA contributed to the interpretation of the data and provided intellectual input. All authors read and approved the final manuscript.

## Competing interests

The authors declare that they have no competing interests.

